# Early-Life Stress Impairs Perception and Neural Encoding of Rapid Signals in the Auditory Pathway

**DOI:** 10.1101/2022.06.14.496208

**Authors:** Yi Ye, Michelle M. Mattingly, Matthew J. Sunthimer, Jennifer D. Gay, Merri J. Rosen

**Affiliations:** Hearing Research Group, Department of Anatomy and Neurobiology, Northeast Ohio Medical University, Rootstown, OH, USA; Brain Health Research Institute, Kent State University, Kent, OH, USA; Department of Otolaryngology, Head and Neck Surgery, Rutgers Robert Wood Johnson Medical School, New Brunswick, NJ, USA

**Keywords:** development, early-life stress, auditory cortex, auditory brainstem response, gap detection, temporal processing, auditory perception, acoustic startle

## Abstract

In children, early ear infections are a risk factor for later speech perception deficits. This is likely because auditory deprivation during a developmental critical period (CP) induces long-lasting deficits in perception and ACx encoding of temporally-varying sounds. CPs also create susceptibility to early-life stress (ELS) in neural regions involved with cognition and anxiety. As CP mechanisms are shared by sensory cortices and higher neural regions, ACx and temporal encoding may also be susceptible to ELS. To examine the effects of ELS on temporal processing, we developed a model of ELS in the Mongolian gerbil, a well-established model for auditory processing. ELS induction impaired the behavioral detection of short gaps in sound, which are critical for speech perception. This was accompanied by reduced neural responses to gaps in ACx, the auditory periphery, and auditory brainstem. These ELS effects presumably degrade the fidelity of sensory representations available to higher regions, and could contribute to ELS-induced problems with cognition.

## Introduction

Sensory experience early in development is essential for the normal maturation of cortical circuits, allowing for accurate perception of the sensory world. In the auditory modality, children who experience even mild to moderate hearing loss over sensitive time windows of development are at risk for later problems with speech perception (Schonweiler et al., 1998; Psarommatis et al., 2001; Whitton and Polley, 2011). This is likely due to auditory deprivation during early postnatal critical periods for neural plasticity. Critical periods are defined developmental windows where the brain undergoes increased susceptibility to alterations by sensory experience (Takesian and Hensch, 2013). Altering auditory experience during these time windows (by manipulations such as inducing partial or complete hearing loss, or exposure to altered environmental soundscapes) causes impairments in auditory perception and alters the sound-evoked response properties of auditory cortical (ACx) neurons (Zhang et al., 2002; Han et al., 2007; Rosen et al., 2012; Gay et al., 2014; Park et al., 2015; Ihlefeld et al., 2016; Green et al., 2017; Yao and Sanes, 2018). In particular, the neural circuitry that encodes temporally-varying signals develops gradually postnatally, and is thus susceptible to auditory deprivation during the critical period for ACx development (Sanes and Woolley, 2011; Mowery et al., 2015). This is important because the ability to detect rapid changes in sound (auditory temporal processing) is intrinsic to deciphering our soundscape, including analyzing auditory scenes and understanding speech (Tallal, 2004). In sensory cortices, critical period plasticity is driven by changes in multiple neural elements, including inhibitory neural maturation, developmental regulators such as neurotrophins, and perineuronal nets (Reh et al., 2020).

Early-life stress (ELS) is well-known to affect critical period mechanisms identified in studies of sensory cortices, but these effects have been observed in higher neural regions rather than sensory cortices. ELS in both humans and animal models can elicit long-term effects on anxiety, memory, learning, and cognition (Cameron et al., 2017). These behavioral changes have been intensely investigated in neural regions underlying those higher-level processes, including the frontal cortex, hippocampus, and amygdala. In those regions, ELS affects the maturation of inhibitory neurons, perineuronal nets, brain-derived neurotrophic factor (BDNF), and dendritic morphology (Bath et al., 2013; Castillo-Gomez et al., 2017; Murthy et al., 2019; Page and Coutellier, 2019). Because these mechanisms underlie sensory cortical plasticity during critical periods (Hensch, 2005), this suggests that ELS may also alter primary cortical sensory regions. However, the effects of ELS on auditory cortical neuronal properties, subcortical auditory regions, or behavioral sensitivity to rapid sounds have not been examined directly.

The literature indicates that early-life stress may affect the auditory system. In rats, ELS reduced the magnitude of auditory evoked potentials (Ellenbroek et al., 2004; Yates et al., 2016). In humans, childhood maltreatment, low socioeconomic status (SES), and low maternal education are proxies for developmental stress. Children raised in these situations have been shown to have altered gray matter volume in ACx and diminished auditory fiber tract integrity (Teicher et al., 2016), lower amplitude auditory brainstem responses (ABRs) (Skoe et al., 2013), and poorer performance on speech perception tasks (Nittrouer and Burton, 2005). Although the underlying mechanisms may differ, stress induced during adulthood alters auditory perceptual sensitivity (Ashton et al., 2000; Hasson et al., 2013; Perez et al., 2013) and alters neural morphology and response properties in ACx and inferior colliculus, the auditory hub of the midbrain (Bose et al., 2010; Dagnino-Subiabre et al., 2012; Ma et al., 2015). Thus, ELS may have widespread effects on the auditory pathway and even the auditory periphery; for example, a corticotropin-releasing factor signaling system is active in the cochlea (Vetter and Yee, 2018).

We examined whether ELS affects auditory temporal processing – specifically, the detection of short gaps in continuous background sound. This ability can be psychophysically quantified by measuring detection thresholds for brief gaps in sound (gap detection thresholds, GDTs). Gap detection represents a general measure of temporal acuity and requires an intact auditory cortex (Green, 1971; Ison et al., 1991; Threlkeld et al., 2008). Because the cortical circuitry that encodes detection of short gaps develops gradually postnatally, it may be affected by early-life stress, similarly to how this circuitry is affected by early auditory deprivation (Rosen et al., 2012; Mowery et al., 2015; Green et al., 2017).

Here, we induced early-life stress in Mongolian gerbils, an established animal model for auditory processing, in a time window encompassing the critical period for ACx maturation. ELS gerbils showed poorer behavioral gap detection, and the responses to short gaps in sound were reduced in ACx neurons and in the auditory nerve and brainstem. Because gap detection in children is predictive of later speech processing abilities (Benasich et al., 2006; Muluk et al., 2011), ELS-induced deficits in gap detection implicate ELS as a risk factor for persistent problems with speech processing.

## Materials & Methods

### Subjects and Experimental Design

All procedures relating to the maintenance and use of animals were approved by the Institutional Animal Care and Use Committee at Northeast Ohio Medical University. Mongolian gerbils (*Meriones unguiculatus*) from multiple litters were housed with littermates in a 12 hour light/dark cycle. Control and ELS animals were from separate litters, to avoid any stress on Control animals or parents induced by removing siblings for ELS treatment. The numbers of animals used for each measurement are indicated in the Results.

The experimental design is shown in Figure 1A. ELS animals experienced stress induction from P (postnatal day) 9-24 as described below, while Control animals were left undisturbed in their home cages. Bodyweight was measured from a subset of animals on days of stress induction. Blood samples were collected from a subset of animals at P25 (the day following the last stress induction event) or P33 (immediately following one session of gap-PPI) to measure corticosterone levels. Auditory brainstem responses were collected from a subset of animals within an age range of P33-40. Animals used for bodyweight, corticosterone, or ABRs did not contribute to any other measurements.

**Figure 1:**
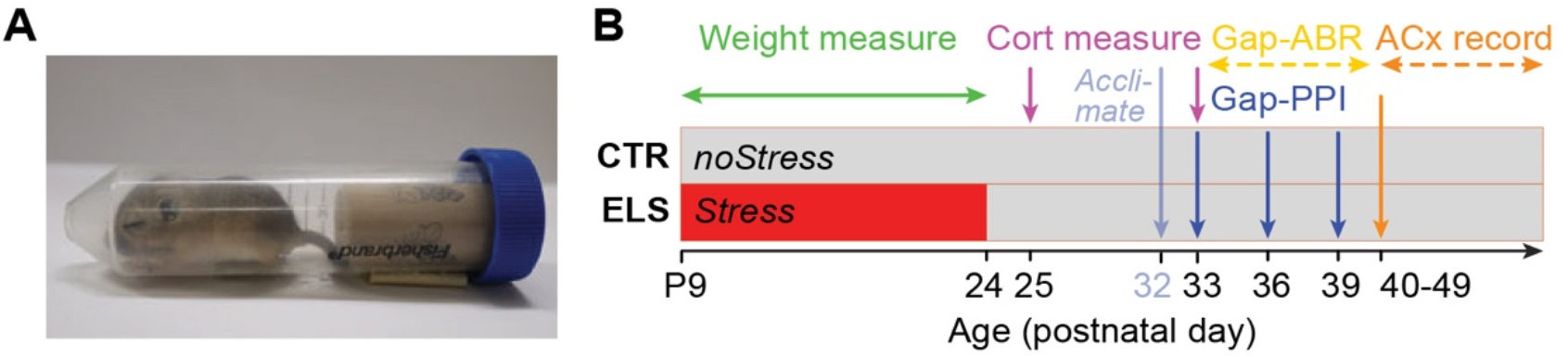
Experimental design and early-life stress induction method. A) ELS was induced by intermittent maternal separation and restraint as depicted, placing the restrained animals in isolated enclosures for ten 2-hour sessions at unpredictable times over the age range P9-24. B) The experimental design timeline depicts measures of auditory processing of gaps in ongoing sound (gap-PPI for behavioral sensitivity, gap-ABR for peripheral and brainstem sensitivity, and ACx recordings for cortical sensitivity), along with measures verifying successful stress induction (bodyweight, blood corticosterone, and startle-only amplitude collected during gap-PPI sessions).

On P33, P36, and P39 (following one hour of acclimation to the startle testing enclosure on P32), a subset of animals were behaviorally tested in one-hour acoustic startle gap-PPI sessions (the behavioral setup allows testing of 8 animals simultaneously). Over the next ∼14 days (P40 – P54), these animals underwent cortical recordings (one animal per day). Animals for cortical neurophysiology were chosen from each group without regard to behavioral performance. Each animal yielded a variable number of neurons for analysis, so recordings were conducted until a sufficient number was collected across groups (see Results).

### Early-life stress induction

ELS was induced postnatally by a combination of maternal separation and restraint from P9-24. Ten two-hour restraint sessions occurred either in the morning or late afternoon within this 16-day time window, to reduce predictability of the restraint session; unpredictability is widely known to be a strong contributor to stress-related behavioral changes (Willner, 1997; Mineur et al., 2006; Bath et al., 2017). During these sessions, the home cage was transported from the animal facility to the testing room, and pups were separated from their parents and littermates and individually isolated. P9 – P24 animals were placed into 50ml centrifuge tubes. The tubes were perforated to allow for easy breathing, and were partly filled with a cork to prevent the animal from free movement. The tubes were placed either horizontally or vertically each in their own small, sound attenuated, anechoic booth, with the lights on. Animals that were too large for the centrifuge tubes (≥ P20) were placed in a small restrainer, and then placed in the booth. The booths and small restrainers were those used for later behavioral acoustic startle testing to measure gap detection abilities. After each session, pups were returned to their home cages and then to the animal facility. ELS was induced in all pups of a litter. While this precludes using siblings as controls, it importantly prevents stress being unintentionally induced in control animals or parents when the home cage and pups are regularly disturbed and removed.

### Corticosterone measurement

To measure corticosterone levels, blood was collected from the retro-orbital vein in animals briefly anesthetized with isoflurane. Blood was centrifuged at 3000 rpm for 30 min, and plasma was separated and saved at -80°C. Plasma corticosterone levels were assessed by Enzyme-Linked Immunosorbant Assay (ELISA) kits (Enzo Life Sciences). Plate readings were obtained with a SynergyTM 4 spectrophotometer (BioTek Instruments, Inc.) at a wavelength of 405 nm. Raw optical density was read out between 570 and 590 nm and corrected with a blank control. The blood corticosteroid level was calculated by fitting the raw data with a 4-parameter logistic curve.

### Behavioral testing for gap detection

Gap detection abilities were assessed as inhibition of the reflexive acoustic startle response, where a detectable change that precedes a startling sound inhibits the startle (known as pre-pulse inhibition). The strength of inhibition corresponds with an animal’s detection of the change. Here, the change was a silent gap in background noise (a variation on pre-pulse inhibition, gap-PPI). The procedure has been described previously (Green et al., 2016). Briefly, animals were placed inside a small acoustically transparent restrainer, on a force plate in a sound attenuated, anechoic booth, with the lights on. Two separate speakers in each booth presented either background noise at 50 dB SPL or a startling stimulus at 110 dB SPL (Kinder Scientific Inc., Poway CA). The background noise was bandpassed from 2.5 – 20kHz. We presented 190 trials in pseudorandom order. Of these, 46 trials were startle–only, with a startle stimulus of 20 ms broadband noise at 110 dB SPL, 1 ms rise/fall time. The remaining 144 were gap trials, where the startle stimulus was preceded (by 50ms) by a silent gap in the noise background of either 2, 3, 5, 7, 10, 25, 50, or 125 ms, with 18 trials of each gap duration. The background noise preceding and following the gap was shaped with a 1ms rise/fall time. At the beginning of each session, 5 startle-only trials were presented (not included in analysis) to habituate the responses to a steady-state level (Ison et al., 1973). Sessions lasted one hour.

### Behavioral data analysis

A gap detection threshold (GDT) was calculated for each animal at each session using custom MATLAB scripts (MATLAB, The Mathworks, Natick MA; D. Green and M. Rosen), as described previously (Longenecker et al., 2016; Green et al., 2017). First, the peak response magnitude to the startle stimulus was measured in the time window 20–50 ms after startle stimulus onset. Because the distribution of peak responses had a strong positive skew, a log10 transform was applied to generate a normal distribution of responses within each trial type. Then we determined the peak response threshold at which a reduction in startle was considered statistically significant. To do so, the median values of the peak log-transformed responses for startle only and each gap duration were plotted, and a cubic spline was fitted to this plot, creating a detection function. To find where that function crossed a detection criterion, the transformed startle-only values were bootstrapped (sampled with replacement 10,000 times) to generate a normal distribution, from which 95% confidence intervals were calculated. The lower confidence interval was the value where a reduction in startle indicated significant detection (Fechter et al., 1988). GDT was the shortest gap duration at which the fitted detection function crossed the lower confidence interval. Group differences across sessions were assessed with mixed ANOVAs.

### Gap-ABRs (Auditory Brainstem Responses)

Animals were anesthetized with ketamine (75 mg/kg) and xylazine (5 mg/kg) and presented with auditory stimuli (RZ6 Auditory Processor, BioSigRP software, Tucker Davis Technologies (TDT), Alachua FL). For a subset of animals, behavioral GDTs were measured prior to ABR GDTs; these animals were anesthetized with dexdomitor during ABR recordings (0.2 mg/kg). Responses to individual stimuli were conducted using stainless steel needle electrodes inserted subdermally at the dorsal midline between the eyes (non inverting), posterior to the right pinna (inverting), and base of the tail (common ground), amplified (20x, TDT low-impedance RA4LI), bandpass filtered (0.3–3 kHz), and digitized (24.4kHz, RZ5 BioAmp Processor, TDT). Auditory stimuli were presented at 40 dB SPL from a freefield speaker located 7 cm from the right ear. Stimuli were two bursts of 50 ms bandpass noise (2.5 – 20 kHz) with 1 ms rise/fall times for each burst. The bursts were separated by gaps of varying durations to match those used in behavioral gap detection testing: 2, 3, 5, 7, 10, 25, 50, or 125 ms (Figure 4B). Burst pairs were repeated at intervals of 1 second. Responses were averaged over 200 presentations.

Each wave of the ABR was quantified separately (Figure 4A). Gerbil wave i (equivalent to human wave I) is understood to arise from the distal portion of the auditory nerve (Achor and Starr, 1980; Boettcher, 2002). Gerbil waves ii and iii (equivalent to human wave III) arise primarily from the cochlear nucleus (Moller and Jannetta, 1982; Melcher et al., 1996). Gerbil wave iv (equivalent to human wave V) reflects activity from the cochlear nucleus, superior olivary complex, and inputs to the inferior colliculus (i.e., lateral lemniscus) (Hashimoto et al., 1981; Moller and Jannetta, 1982, 1983). Amplitudes and latencies were measured for each wave, in response to both noise bursts. Peak amplitudes were measured as the voltage difference between each wave peak and the following trough, allowing calculation of the amplitude ratio: attenuation of the 2^nd^ burst normalized by the response to the 1^st^ burst. Peak latencies were measured as the latency to each peak from the start of each noise burst. This allowed calculation of any latency shift: the difference in the response latency to the 2^nd^ burst verus the response latency to the 1^st^ burst. Detection threshold for each animal’s gap-ABR was determined as the shortest gap duration with a visible response to the 2^nd^ noise burst in any wave, though typically waves ii or iii were the last waves to disappear as gaps shortened. Group differences across gap durations were assessed with mixed ANOVAs.

### Surgical Preparation

Gerbils were prepared for auditory cortical recordings as previously described (Mattingly et al., 2018). Briefly, animals were anesthetized with isoflurane and held in a stereotaxic apparatus. A small headpost was positioned along the midline and secured with dental acrylic, and a silver ground wire was implanted into the posterior contralateral skull. A craniotomy was made over the left temporal cortex caudal to the bregma suture and the dura was left intact. A thin well of dental acrylic was built along the perimeter of the craniotomy, the cortical surface was covered with silicone oil, and the craniotomized area was covered with a disposable cap of silicone elastomer (ImageLB-28, Matrics Inc. Osseo MN). The entire skull was covered with dental acrylic to form a headcap.

### Neurophysiological Recordings

On the day of recording, animals were anesthetized with urethane (1.3g/kg, administered in 2 doses over 1.5 hrs) and placed in a soundproof chamber (Industrial Acoustics Company, North Aurora IL) on a heating pad. The head was stabilized using the headpost, the silicone elastomer cap removed, and the dura was covered with saline during recording to maintain moisture. The dura was nicked and platinum-plated tungsten electrodes (1.5-2.5 MΩ; MicroProbe, Gaithersburg, MD) were advanced to isolate neurons in primary ACx based on response characteristics (reliable, short-latency, non-adapting responses to tones). Electrical signals from the brain were amplified (250x, TDT RA16PA Medusa preamplifier), filtered (0.25 to 10 kHz), and digitized (24.4 kHz, TDT RZ5 BioAmp Processor). The TDT equipment was controlled by custom software written in MATLAB and TDT RPvdsEx programming environments (TytoLogy by S.J. Shanbhag). Units were isolated by spike amplitude. Spikes were detected offline (Plexon Offline Sorter, Dallas TX), and sorted based on spike shape and principal component analysis.

### Acoustic Stimulation for Neurophysiology

TDT equipment (RZ6 Auditory Processor) was used to deliver auditory stimuli and record neural responses using custom software written in MATLAB and TDT RPvdsEx (TytoLogy by S.J. Shanbhag; modified by M.J. Rosen). Auditory stimuli over a frequency range of 200Hz – 35kHz were calibrated (custom MATLAB scripts, S.J. Shanbhag). Calibration data were collected using a ¼ inch microphone (Brüel and Kjær (B&K) model 4939), a preamplifier (B&K 2670) and a conditioning amplifier (B&K Nexus model 2690). Stimuli were delivered through a freefield speaker positioned 25cm in front of the animal.

Each ACx unit’s response to tones was assessed by presenting 200ms tone pips (1 sec intertrial intervals, 5 ms cosine-ramped rise/fall). First, the frequency range over which the neuron was responsive was obtained with an iso-intensity function at 60 dB SPL to determine the best frequency (BF) of the unit (the frequency eliciting the highest firing rate). This was followed by a rate-level function (RLF) at BF, measured at increments of 10 dB SPL, for 15 trials. Threshold was visually determined as 5dB below the lowest level with a clear increase in firing above lower levels. A gap detection function was obtained at the BF of the unit and 30dB above threshold, with 20 trials at each gap duration, presented in random order. Gap detection stimuli consisted of two consecutive tone bursts lasting a total of 400ms (with 5 ms cosine-ramped rise/fall). The bursts were separated by gaps of varying durations matching those presented behaviorally (0, 1, 2, 3, 5, 7, 10, 15, 25, 50, or 125 ms with a 0.5 ms cosine-ramped rise/fall) inserted between the two bursts at 200ms after the onset of the first burst. The 0 ms gap stimulus was the control against which gap detection was measured, and thus did not contain rise/fall ramps at 200ms, but was instead a continuous 400ms tone burst. This control was chosen to mimic the behavioral stimuli, where detection of gaps in background noise were calculated in comparison with continuity in the background noise. After collecting the gap detection tone function, the unit was presented with 200ms bursts of noise at 50dB SPL, bandpassed from 2.5 – 20kHz, to match the background noise used for the behavioral testing. If the unit showed a clear response to the noise, a noise gap detection function was obtained with 20 trials at each gap duration, presented in random order.

### Neural Data Analysis

#### Rate-level functions (RLF)

Firing rates to tones were calculated over a time window equal to the stimulus duration (200ms). Threshold, dynamic range, and monotonicity were determined from the RLF. Threshold was defined as the dB SPL level at which there was at least a 35% increase in firing rate, stepping up from one level to the next; threshold firing rates were calculated at this sound level. Dynamic range was defined as the range between the sound levels where each cell responded at 10% and 90% of its maximum firing rate, calculated by interpolation. A monotonicity index (MI) was calculated by dividing the firing rate at the maximum presented sound intensity (80 dB SPL) by the maximum firing rate within the RLF (Moore and Wehr, 2013). The MI ranges from 1 (no intensity tuning) to 0 (strong intensity tuning). All data were analyzed with custom MATLAB scripts (M.J. Rosen). Group differences were assessed with Kruskal-Wallis (KW) nonparametric ANOVAs.

#### Gap detection

Responses to each gap duration were measured from post-stimulus time histograms (PSTHs) with 5ms bins calculated across the 20 trials. Only units with onset responses to the first burst of the gap detection stimuli were used for gap detection analysis. A valid onset response (within 100ms following the first burst) was determined as spiking 20% of the time within a single bin across trials. A valid gap response (within 100ms following gap offset (equivalent to second burst onset)) was determined based on the bin with maximal firing in this window. We evaluated the response following gap offset (rather than gap onset (equivalent to offset of the first burst)) because 1) most units did not have clear offset responses, and 2) cortical responses at gap offset have been directly linked to behavioral gap detection (Weible et al., 2014). After subtracting baseline firing (calculated in the same window when no gap was present), a response in the peak bin 20% of the time was considered a valid gap detection response. Experimenters blinded to animal treatment then visually inspected PSTHs across all gap durations to verify that the detected responses following the second burst were consistent across gap durations (as in Figure 5A-D). This effectively excluded spurious responses that could arise with use of a 100ms evaluation window. The shortest duration gap with a valid gap detection response was considered the GDT (Eggermont, 2000). If a significant response to the second burst was absent for all gap durations, a GDT of 135ms was assigned (since only cells with valid onset responses were analyzed, a gap longer than the longest presented (125ms) would necessarily elicit a response). First spike latency (FSL) and FSL jitter were measured directly from spike timing rather than from binned PSTHs, and were based on the onset response to the first burst. All data were analyzed with custom MATLAB scripts (M.J. Rosen). Group differences were assessed with Kruskal-Wallis nonparametric ANOVAs.

#### Ideal observer analysis

An ideal observer classifier model was used to determine how well the population of neural units could discriminate between gap and no-gap trials. By definition, an ideal unbiased observer maximizes hits and minimizes false alarm rates to perform a given task optimally (e.g., (Geisler, 2011). We compared the ideal observer’s ability to detect gaps of varying durations. We trained a support vector machine with 85% of the neural data, and ran a 10-fold cross-validation test of classifier performance with the remaining 15% of the neural data, separately for Control and ELS groups. The classifier was trained and tested with trial-by-trial firing rates from 100ms time windows following gap offset. The classifier, implemented in MATLAB 2015a, used custom scripts (M.J. Rosen, Y.E. Cohen & A. Ihlefeld) that trained a binary support vector machine classifier (fitcsvm in MATLAB) and tested the performance on the remainder of the data by computing a loss estimate using cross validation (crossval in MATLAB). Equal numbers of neurons were used across the treatment groups, limited by the group with the fewest units that had valid onset responses to the first burst (104 units for trials with tone carriers at BF; 94 units for trials with bandpass noise carriers). The cross-validated classifier model was run 100 times to generate mean performance with SEM error bars.

## Results

### Establishing a model of early-life stress in the Mongolian gerbil

While early-life stress is frequently studied in rodent models, to date there is not an established model for ELS in the gerbil. Here, we induced early-life stress through a combination of intermittent, unpredictable maternal separation and restraint over the developmental window P9-24 (Figure 1A and B). This time window encompasses the critical period for the maturation of several auditory cortical properties, including evoked firing rates (Mowery et al., 2015). Furthermore, auditory deprivation during this time window is known to induce perceptual deficits (Caras and Sanes, 2015; Green et al., 2017). Thus, induction of stress over this time window has the potential to affect development of the auditory cortex. To assess the effectiveness of the stress induction, we measured three established physiological markers of early-life stress: blood corticosterone levels, bodyweight gain over development, and response to a startling stimulus (Buynitsky and Mostofsky, 2009; van Bodegom et al., 2017; Demaestri et al., 2020).

Stress enhances the activity of the hypothalamus-pituitary-adrenal (HPA) axis, resulting in increased secretion of corticosteroids from the adrenal cortex of the adrenal gland (van Bodegom et al., 2017). In rodents including gerbils, plasma corticosterone (CORT) is the main glucocorticoid involved in regulating stress responses, and is thus often used as a biomarker for stress. Chronic or developmental stress can either increase or decrease CORT levels, depending on the details of stress induction. To establish whether our ELS induction protocol affected baseline levels of this stress hormone, we measured corticosterone levels in blood samples collected from animals at P25, the day following the last stress induction event (Figure 1A). To determine whether ELS induction affected CORT levels following an acute stressor, we collected blood samples at P33 in a separate group of animals, immediately following a session of acoustic startle gap-PPI measurements. CORT levels were significantly lower in ELS than Control animals, both at baseline and following this acute stressor (Figure 2A; Control: baseline mean CORT (n = 19) = 11570.0 pg/ml; acute mean CORT (n = 18) = 441.5 pg/ml; ELS: baseline mean CORT (n = 20) = 3815.0 pg/ml, acute mean CORT (n = 20) = 326.0 pg/ml; one-way ANOVAs: baseline F_(1,37)_ = 17.53, p < 0.001, Ƞ^2^ = .321; acute F_(1,36)_ = 4.197, p = 0.048, Ƞ2 = .104). These data demonstrate that the P9-P24 ELS induction protocol affects the HPA axis in Mongolian gerbils.

**Figure 2:**
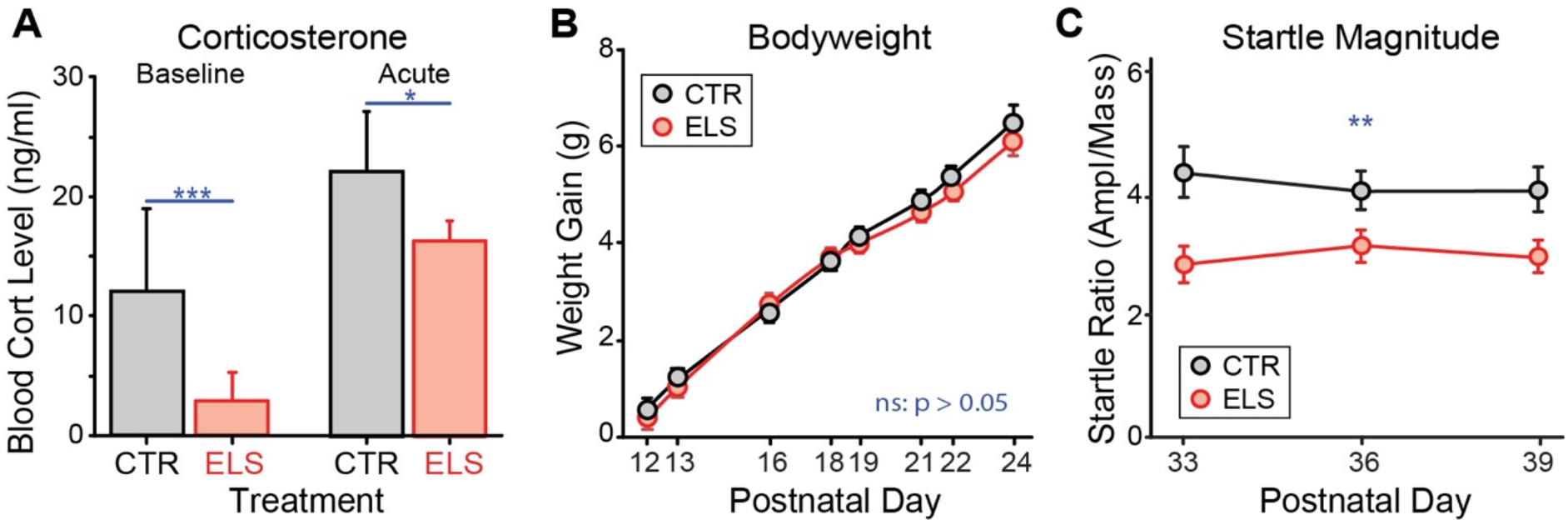
Successful induction of early-life stress in the gerbil, without a nutritional deficit. A) Blood serum corticosterone levels were reduced by ELS in two behavioral situations: baseline (P25, one day after the completion of ELS induction) and following an acute stressor (P33, immediately after first experiencing a gap-PPI acoustic startle test session). B) Bodyweight gain over postnatal development was not affected by ELS induction. C) Startle reactivity was reduced in ELS animals. *** p ≤ 0.001; ** p ≤ 0.01; * p < 0.05.

When rodents are raised with ELS, their body weights may be reduced compared with Control animals, though this varies depending on how ELS is induced (Demaestri et al., 2020). We weighed ELS gerbils after each restraint session (at postnatal days 12, 13, 16, 18, 19, 21, 22 and 24), and weighed Control animals at the same ages. Body weight gain did not differ across this developmental period (Figure 2B, repeated measures ANOVA: F_(1,24)_ = 1.159, p = 0.29, n = 10 Control, n = 16 ELS). This reduces the likelihood of differential nutritional consequences, or of any potential confound arising from comparing animals of different body weights in a startle paradigm.

Startle reactivity measured by the acoustic startle response, which involves a simple subcortical circuit, can be modified by hormones of the HPA axis (Lee and Davis, 1997). To determine whether our ELS induction protocol altered startle reactivity, we measured the amplitude of startle-only responses (normalized by each animal’s mass) during behavioral testing for gap detection using gap-PPI. The normalized amplitudes of startle responses in trials that did not contain gaps were compared between Control and ELS groups, for each of three sessions of gap-PPI testing. A mixed ANOVA with a between-subject factor of treatment and a within-subject factor of session revealed a significant effect of treatment, with lower startle-only amplitudes in ELS animals (Figure 2C, F_(47,1)_ = 7.734, p = 0.008, Ƞ^2^ = .141, n = 26 Controls, n = 23 ELS). This reduced startle reactivity is consistent with the lower circulating levels of corticosterone in the ELS gerbils.

### Behavioral gap detection is impaired by ELS

Having established that our developmental manipulation effectively induces early-life stress, we measured behavioral auditory sensitivity to temporally-varying sounds that require auditory cortex: brief gaps in an ongoing noise (Ison et al., 1991; Kelly et al., 1996; Syka et al., 2002; Threlkeld et al., 2008). We chose to use a reflexive, pre-attentive measure of gap detection: gap-PPI of the acoustic startle response. PPI, a measure of detection rather than perception, is not influenced by attentional factors provided the interval between stimulus and startle is ≤ 60 ms (Li et al., 2009), but see (Cope et al., 2022). This allows measurement of auditory processing without the potentially confounding contribution of cognition or attention, both of which can be affected by stress.

The day before testing, animals were acclimated to the testing enclosure for 1 hour. Testing sessions were conducted at the same time each day, on P33, 36, and 39, and GDTs were calculated for each session (Figure 1A; animals tested for GDTs were not used for body weight or CORT measures). A mixed ANOVA with a between-subject factor of treatment and a within-subjects factor of test session revealed significantly poorer (higher) GDTs for ELS animals, collapsed across sessions (Figure 3A; F_(1,47)_ = 7.105, p = 0.011, partial Ƞ^2^ = .131; n=26 Control, n = 23 ELS). There was also greater variability across the ELS animals (Levene’s test F_(1, 35)_ = 7.59, p < 0.0001). Posthoc tests with Sidak corrections for multiple comparisons showed that ELS GDTs were worse than Controls in sessions 2 and 3 (p = 0.005 and 0.014), though not in session 1 (p = 0.56). Even when examining the best performance for each animal, ELS animals had poorer and more variable thresholds than Controls (Figure 3B, t_(44)_ = -2.506, p = 0.008; Levene’s test: F_(1, 47)_ = 9.94, p = 0.002). Beyond examining just threshold, the response ratio comparing gap to no-gap startle amplitudes depicts sensitivity across gap durations, with smaller amplitude indicating better detection. The Controls had smaller amplitude ratios across gap durations than ELS (Figure 3C, main effect of treatment: F_(1, 47)_ = 232.7, p < 0.001, partial Ƞ^2^ = .83), with group differences at longer durations for session 1 (50 and 100ms, p = 0.026 and 0.046), but shorter durations for sessions 2 and 3 (7, 10 and 25ms: session 2, p = 0.026, 0.009, 0.002; session 3, p = 0.029, < 0.001, = 0.023).

**Figure 3:**
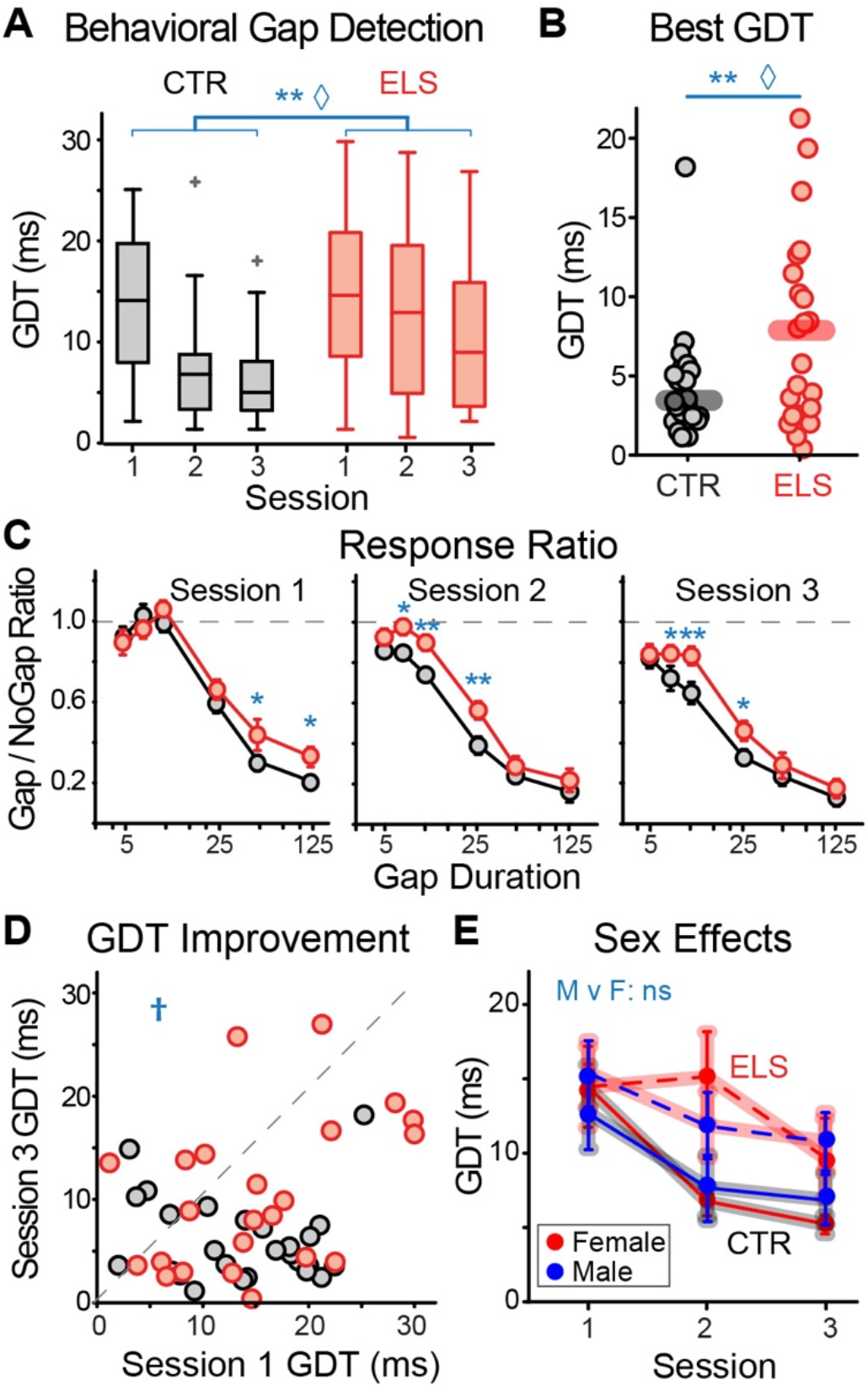
Behavioral gap detection was impaired by early-life stress. A) Across three sessions of gap-PPI, ELS animals had poorer gap detection thresholds, concurrent with greater variability across ELS than Control animals. B) The best threshold across all sessions was worse for ELS animals. C) Gap to no-gap response ratios were higher for ELS animals, indicating poorer detection overall. Controls showed greater improvement based on response ratio across sessions than ELS. D) For gap detection thresholds, there was a trend for Controls to improve more across gap-PPI sessions than ELS animals. E) Males and females performed equivalently to one another within each treatment group. *** p ≤ 0.001; ** p ≤ 0.01; * p < 0.05; † p < 0.1; ◊: unequal variance, p < 0.005; in boxplots, box edges are 25^th^ and 75^th^ percentiles, with whiskers extending to the most extreme data points excluding outliers.

Testing across multiple days allowed us to assess learning, indicated by improvement of gap detection. Both groups of animals improved their GDTs across sessions (sessions 1 vs 3: CTR: p < 0.0001, ELS: P = 0.009). While not significant, there was a trend for CTR animals to improve thresholds across sessions more than ELS animals (Figure 3A and D, sessions 3 vs 1: t_(47)_ = 1.3717, p = 0.088). Consistent with this trend, response amplitudes across all gap durations improved more for Control than ELS animals (comparing sessions 1 and 3, t_(326)_ = 2.45, p = 0.015).

Because ELS often has differential effects on males and females, we applied a mixed ANOVA with between-subject factors of sex and treatment and a within-subject factor of test session. There was no significant effect of sex on GDTs, indicating that ELS affected gap detection similarly in males and females (Figure 3E; ELS n = 10 females, 13 males; Control n = 15 females, 11 males).

### Neural gap detection in the auditory periphery and brainstem is impaired by ELS

The detection of brief gaps in sound (< 100 ms) using the gap-PPI measure is known to require an intact auditory cortex (Ison et al., 1991; Kelly et al., 1996; Syka et al., 2002; Threlkeld et al., 2008), and gap detection is worsened by manipulation of neurons in ACx (Weible et al., 2014). Yet disruption of signal encoding earlier in the auditory pathway could also impact gap detection. We therefore measured the auditory brainstem response to gaps of varying durations, matching those used in the behavioral experiments (Figure 4B). Amplitudes and latencies of the responses to two sequential noise bursts separated by varying gaps were measured for ABR waves i, ii, iii, and iv (Figure 4A). Gap-ABR GDTs (defined as the shortest gap where any response was visible to the 2^nd^ burst) were higher in ELS animals (Figure 4C; one-way ANOVA between CTR (n = 19) and ELS (n = 20): F_(1, 38)_ = 9.882, p = 0.003, Ƞ^2^ = .211). There was also greater variability of GDTs across the ELS animals (Levene’s test: F_(1, 37)_ = 12.86, p = 0.001). This may have been due to a floor effect for the CTR animals, as 70% and 95% of Controls had visible responses at 2 and 3 ms respectively, compared with 26% and 53% of ELS animals.

**Figure 4:**
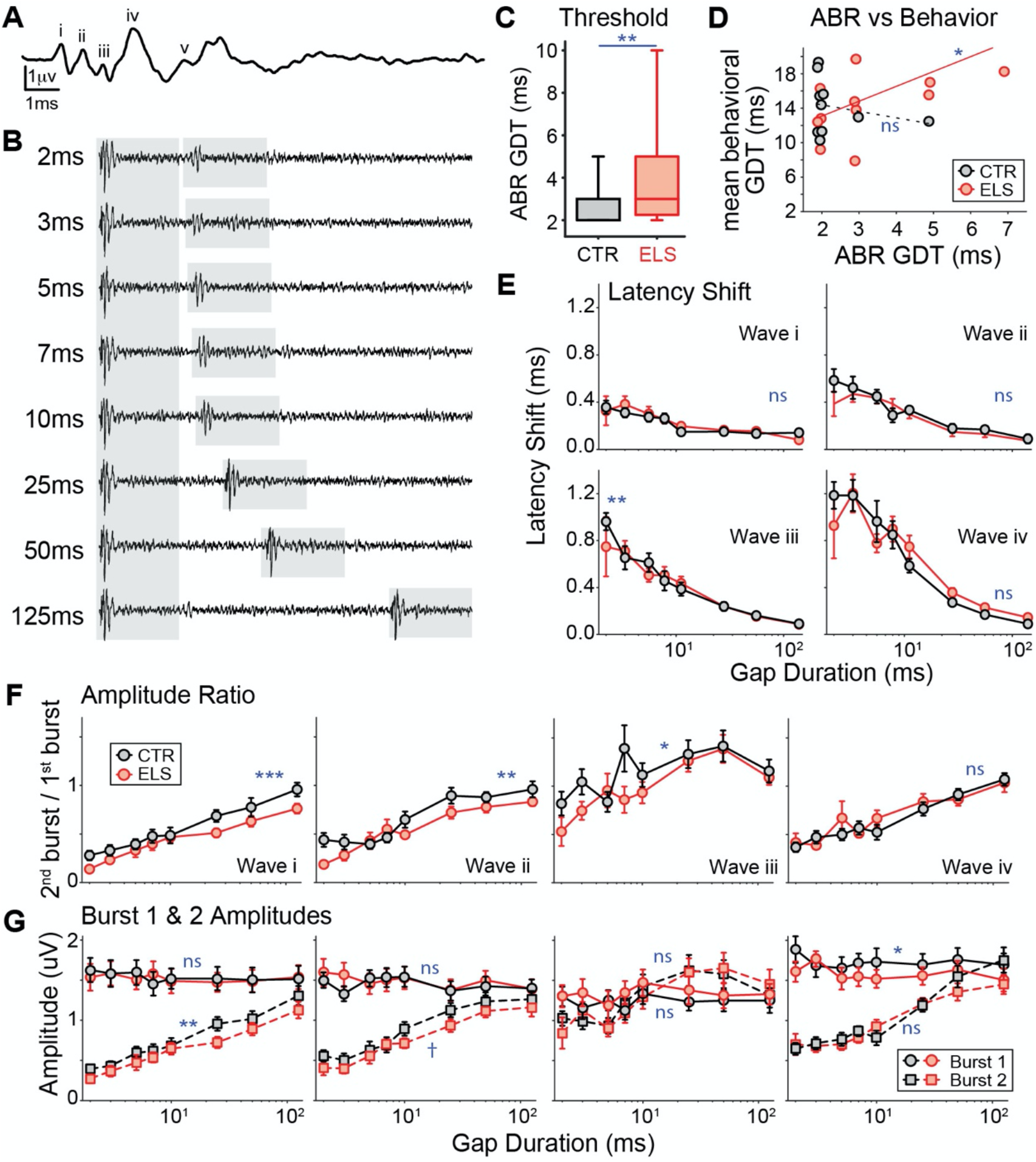
Peripheral and brainstem sensitivity to gaps was reduced by early-life stress. A) Example ABR waveform from gerbil, showing waves i, ii, iii, iv, and v. B) Example ABR response to two 50 ms noise bursts (*gray shaded regions*) separated by gaps of varying durations. C) Higher ABR gap detection thresholds for ELS animals, based on the shortest gap duration with a visible response to the 2^nd^ noise burst. D) For ELS animals, higher gap-ABR GDTs correlated with higher behavioral GDTs, while CTR animals had a floor effect for gap-ABR GDTs. E) ELS did not alter latency shift between the two noise bursts except at the shortest gap duration for wave iii. F) ELS reduced the gap-ABR amplitudes to the 2^nd^ burst normalized by the 1^st^ burst, for waves i, ii, and iii. G) ELS reduced the amplitude to the 2^nd^ burst for waves i and ii, but increased the amplitude to the 1^st^ burst for wave iv. *** p ≤ 0.001; ** p ≤ 0.01; * p < 0.05; † p < 0.1; in boxplots, box edges are 25^th^ and 75^th^ percentiles, with whiskers extending to the most extreme data points excluding outliers.

For a subset of animals, prior to ABR recordings, behavioral GDTs were measured and averaged across the three sessions, to assess any correlation between behavioral and ABR GDTs. Control animals did not show any correlation, likely due to a floor effect for ABR GDTs (Figure 4D; CTR (n = 10): rho = -0.18, p = 0.62). Yet for ELS animals, poorer behavioral GDTs were positively correlated with poorer ABR GDTs (ELS (n = 12): rho = 0.62, p = 0.031). This suggests that in ELS animals, a response following the gap does not occur reliably at the level of the auditory nerve, and thus may not provide high-fidelity signal input to auditory cortex, which is required for the detection of short gaps.

As expected, for both Control and ELS animals and all ABR waves, the response to the 2^nd^ noise burst occurred at a delayed latency compared with the response to the 1^st^ burst, with larger delays following shorter gaps (Figure 4E). Similarly, the response amplitude to the 2^nd^ burst was reduced compared with that to the 1^st^ burst, particularly for shorter gaps (Figure 4F, G). The latency shift did not differ between CTR and ELS animals for ABR waves i, iii, or iv. For wave ii, ELS animals had a smaller latency shift that was confined to the 2ms gap duration, though only 21% of the animals showed visible responses and contributed at this gap duration (Figure 4E; univariate mixed ANOVA, with independent variables of treatment and gap durations: wave ii, F_(1,261)_ = 7.348, p = 0.007, partial Ƞ^2^ = .027; posthoc at 2ms, p < 0.001). However, the amplitude ratio of the two responses (effectively normalizing the 2^nd^ burst response for each animal) was smaller for the ELS animals for waves i, ii, and iii (Figure 4F; wave I, F_(1,261)_ = 10.910, p = 0.001, partial Ƞ^2^ = .04; wave ii, F_(1,261)_ = 7.117, p = 0.008, partial Ƞ^2^ = .027; wave iii, F_(1,261)_ = 4.544, p = 0.034, partial Ƞ^2^ = .018). For wave i and a trend for wave ii, this effect was driven by a reduced response to the 2^nd^ burst (Figure 4G, left two panels; wave i: F_(1,261)_ = 9.324, p = 0.002, partial Ƞ^2^ = .034; wave ii: F_(1,261)_ = 2.765, p = 0.098, partial Ƞ^2^ = .010;). In contrast, for wave iv, there was actually an increased response in ELS animals to the 1^st^ burst (Figure 4G, right panel; F_(1,295)_ = 4.409, p = 0.037, partial Ƞ^2^ = .015).

### Neural gap detection in the auditory cortex is impaired by ELS

Although gap-PPI does not involve attention, the detection of short gaps (< 100 ms) requires an intact auditory cortex (Ison et al., 1991; Kelly et al., 1996; Syka et al., 2002; Threlkeld et al., 2008). Thus the behavioral deficit in the ELS animals suggests that cortical sensitivity to gaps may have been impaired by ELS induced during the critical period for ACx maturation. To test this hypothesis, we recorded single- and multi-unit activity (104 CTR and 108 ELS units) from primary ACx with tungsten microelectrodes during the week following the last behavioral session, when animals were P40-49. Gap-detection functions were collected with carriers of either tones at each unit’s best frequency, or bandpass noise matching that used in the behavioral testing, and GDTs were calculated for each unit. Figure 5A-D depicts 4 different gap-responsive units, to represent the variability of response types and to show different GDTs (stars). Gap detection was measured based on firing immediately following gap offset (i.e., the response to the 2^nd^ sound burst).

**Figure 5:**
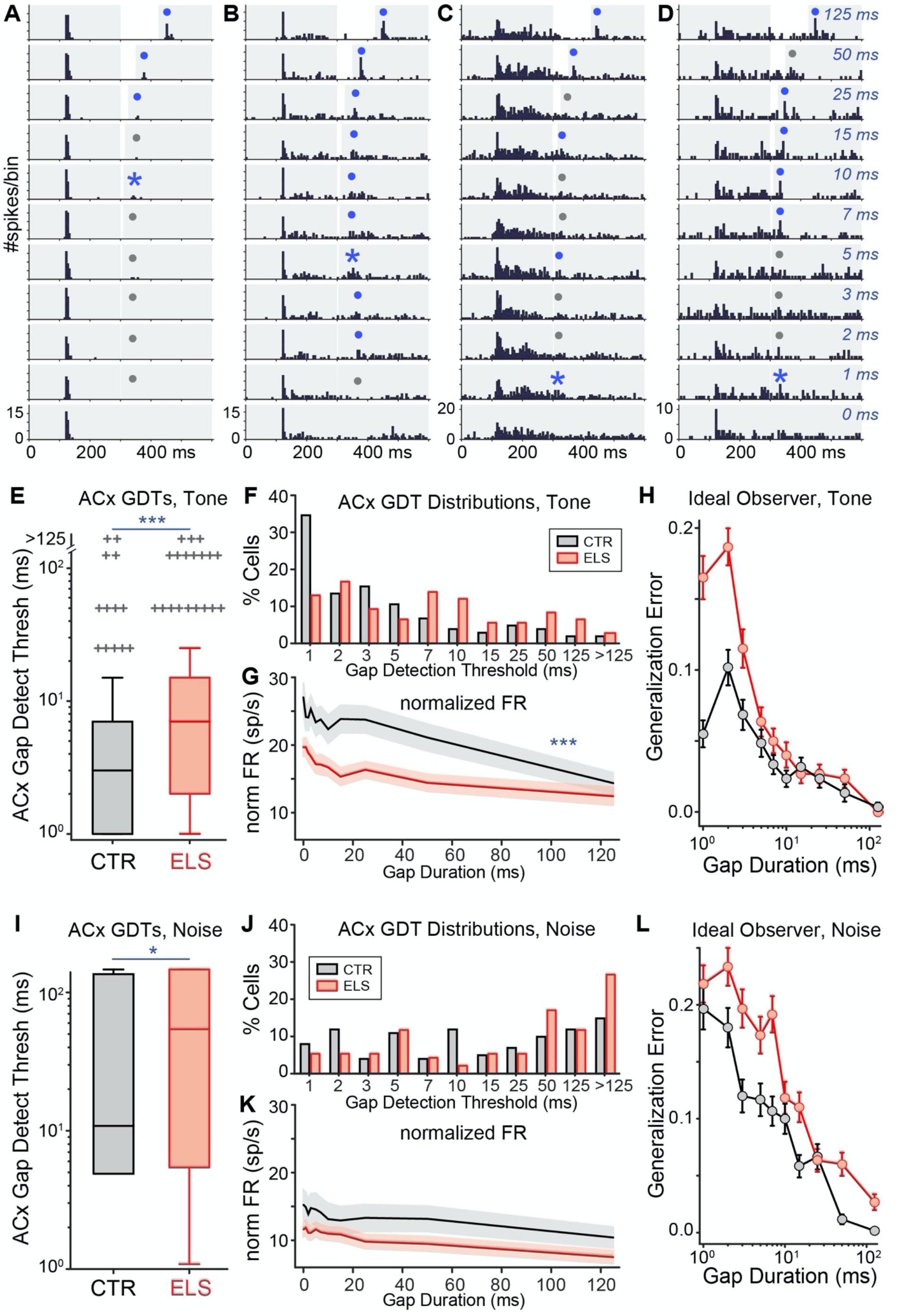
Early-life stress reduced cortical sensitivity to gaps, with a population cortical model predicting poorer behavioral sensitivity. A-D) PSTHs from 4 units in response to gaps of varying durations, where *blue stars* indicate the GDT, and *gray and blue dots* indicate nonsignificant and significant gap responses, respectively. The shortest significant latency was considered the GDT after visual inspection – e.g., in B, the blinded experimenters agreed that the significant responses to 2 and 3ms gaps were at an inconsistent latency compared with the longer gap responses. E) GDTs measured from gap detection functions with tone carriers at the BF of each unit were higher for ELS units. F) Compared with Controls, the distribution GDTs was shifted to the right across ELS units responding to tones, with fewer units sensitive to short gaps. G) Differences in firing rate between the 1^st^ and 2^nd^ burst were larger in Controls, particularly at shorter gap durations. H) A population-based ideal observer model trained with tone-evoked firing rates to gaps of varying durations predicted greater error for short gaps in ELS animals. I) Gap detection functions with a noise carrier matching that used in behavioral testing produced higher GDTs in ELS animals, and higher overall GDTs in both groups than with tone carriers. J) The distribution of noise-evoked GDTs was right-shifted in ELS animals, with more units insensitive to even long gaps. K) Differences in firing rate between the 1^st^ and 2^nd^ burst. L) An ideal observer model trained with noise-evoked firing predicted greater error across all gap durations for ELS animals. *** p ≤ 0.001; * p < 0.05; in boxplots, box edges are 25^th^ and 75^th^ percentiles, with whiskers extending to the most extreme data points excluding outliers; shaded regions around FR indicate SEM.

For tone carriers, cortical GDTs were higher for ELS units than Controls (Figure 5E; Kruskal-Wallis ANOVA: χ^2^_(1,210)_ = 16.8, p < 0.0001). Comparing the distribution of neural GDTs shows a clear shift for ELS animals, with more cells having higher GDTs and fewer cells with sensitivity to short gaps (Figure 5F). Across gap durations, firing following the gap (i.e., in response to the second tone burst) was reduced by ELS, both when normalized by the response to the first tone burst (Figure 5G; χ^2^_(1,2451)_ = 11.1, p < 0.001), and also in response to each tone burst (burst 1: χ^2^_(1,2451)_ = 47.2, p < 0.0001, burst 2 χ^2^_(1,2451)_ = 111.6, p < 0.0001). Gap detection functions collected with a noise carrier matching that used in the behavioral testing (n = 101 Control and 94 ELS units with noise responsivity) showed the same pattern, though GDTs were higher in both groups than with a tone carrier, with many more units insensitive to even 125 ms, the longest gap presented (Figure 5I and J; χ^2^_(1,193)_ = 5.8, p = 0.016). Across gap durations, firing was reduced in response to each noise burst (burst 1: χ^2^_(1,2429)_ = 30.3, p < 0.0001, burst 2 χ^2^_(1,2429)_ = 155.1, p < 0.0001), though normalized firing was not significantly reduced (Figure 5K; χ^2^_(1,2429)_ = 1.2, p = 0.27). Despite the general correspondence between cortical and behavioral GDTs across treatment groups, there was no significant correlation between cortical and behavioral GDTs for either group, using either tone or noise carriers (data not shown).

To determine how well the combined information across the population of neural units could discriminate between gap and no-gap trials, and whether this population discrimination performance predicted behavioral differences across the groups, we trained and tested an ideal observer classifier model using the firing rates following gap offset, i.e., in a time window following the 2^nd^ sound burst. Figure 5G and K depict the means of those firing rates, normalized to the response following the 1^st^ sound burst. Cortical responses at gap offset have been shown to directly influence gap detection measured by gap-PPI (Weible et al., 2014). Thus a model trained with these responses should reflect behavioral GDTs, showing poorer detection at shorter gaps. Furthermore, that model should show poorer performance when trained with ELS neurons.

We compared the ideal observer’s ability to detect gaps of varying durations for both groups, with either tone or noise carriers (Figure 5H and L). Consistent with the behavior, for both Controls and ELS the model predicts greater error distinguishing short gap responses from no-gap responses. Comparing treatments, for tone carriers the model further predicts greater errors detecting short duration gaps (< 5ms) in the ELS group than in Controls. For noise carriers, greater ELS error occurs for both short and long gaps. The model is thus consistent with the poorer behavioral gap detection seen in ELS animals.

### ELS reduces the magnitude of auditory cortical responses

Manipulating auditory experience during the developmental period over which ELS was induced here is known to affect basic response properties of auditory cortical neurons (Zhang et al., 2002; Rosen et al., 2012; Green et al., 2017). We thus assessed whether ELS during that period altered cortical responses to tones. First-spike latencies (FSL) were measured from responses to the first burst of sound in gap detection functions acquired with tone carriers at each unit’s BF. Neither FSL nor FSL jitter were altered by ELS (Figure 6A and B). However, the firing rate evoked by the first sound burst was reduced in ELS animals, as was the spontaneous activity in the 100 ms preceding the first burst (Figure 6C and D; KW ANOVAs, spontaneous activity: χ^2^_(1,221)_ = 19.11, p < 0.0001; sound-evoked activity: χ^2^_(1,221)_ = 3.94, p = 0.047). This effect extended to measures acquired from rate level functions acquired at BF. ELS animals had lower firing rates at sound-evoked threshold, and those thresholds were higher for ELS animals (Figure 6E and F; threshold FR: χ^2^_(1,199)_ = 7.85, p = 0.005; threshold dB: χ^2^_(1,199)_ = 8.66, p = 0.003). The dynamic range was not affected by ELS, but there were fewer non-monotonic units in ELS animals, indicated by a higher monotonicity index (Figure 6G and H; monotonicity index: χ^2^_(1,199)_ = 6.61, p = 0.010).

**Figure 6:**
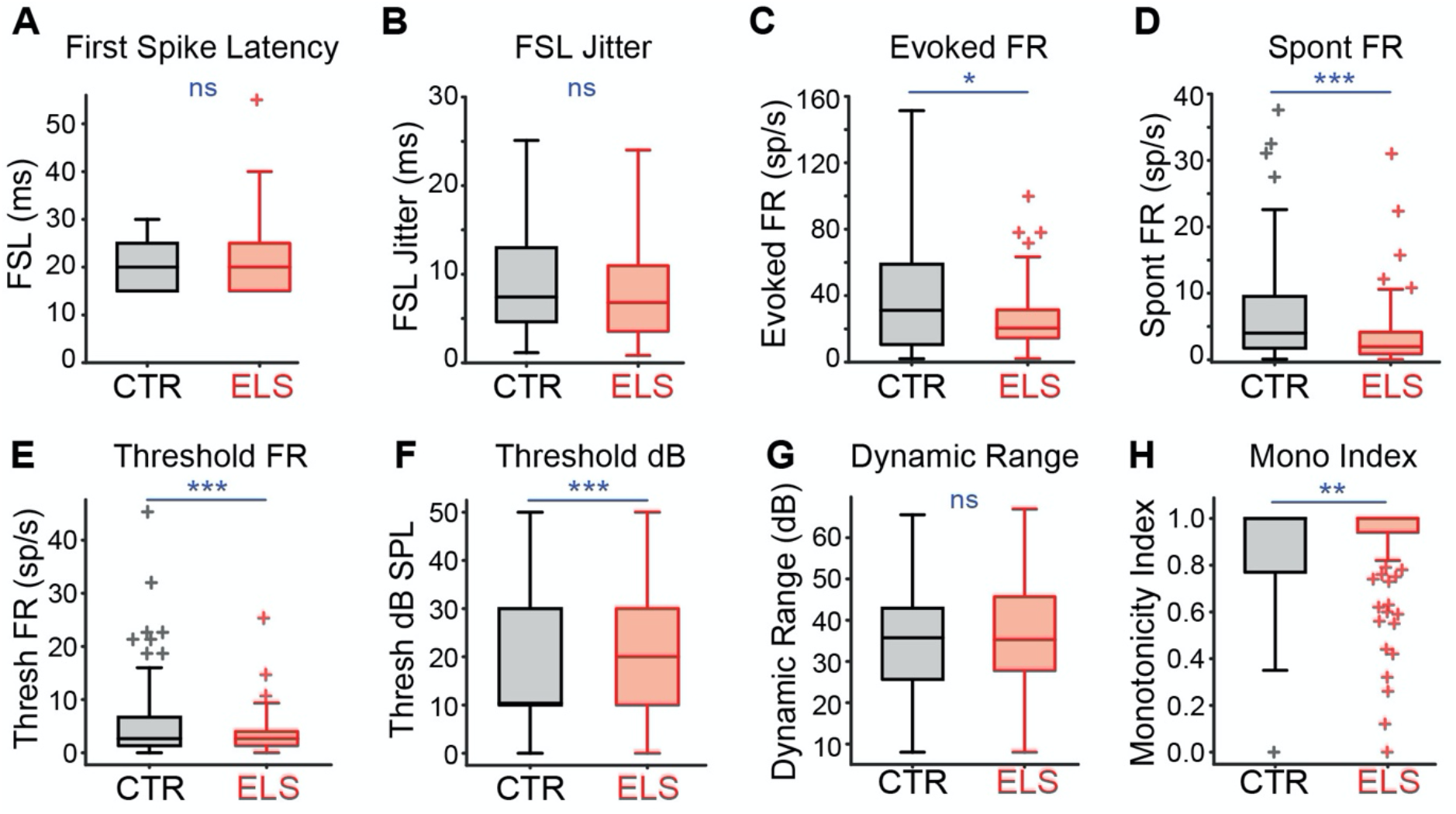
Sound-evoked and spontaneous cortical firing, but not first-spike latency, were reduced by early-life stress. Neither A) FSL nor B) FSL jitter were affected by ELS. In contrast, ELS reduced C) evoked firing, D) spontaneous firing, and E) firing at threshold, along with causing F) higher sound-evoked thresholds. G) Dynamic range was unaffected by ELS, but H) monotonicity was increased by ELS. *** p ≤ 0.001; ** p ≤ 0.01; * p < 0.05; in boxplots, box edges are 25^th^ and 75^th^ percentiles, with whiskers extending to the most extreme data points excluding outliers.

## Discussion

To examine the effects of postnatal stress on auditory temporal processing, we evaluated perceptual and neural gap detection, a common measure of auditory temporal acuity important for the perception of time-varying sounds such as communication calls and speech. We developed a model of early-life stress in the Mongolian gerbil, which was verified by induced shifts in both corticosterone levels and responses to startling stimuli. With this model, we demonstrated that ELS impaired auditory perception of short gaps in sounds, equivalently for males and females. ELS also reduced the improvement in gap detection across behavioral sessions, indicating a learning deficit. Consistent with the necessity of auditory cortex for short gap detection, behavioral deficits coincided with poorer ACx gap detection thresholds and fewer neurons sensitive to short gaps in ELS animals. Further, ABR responses to gaps were reduced for waves corresponding to the auditory nerve and auditory brainstem. Behavioral and ABR gap detection thresholds were correlated in ELS animals, indicating an unreliable response to short gaps at the level of the auditory nerve, and a poor-fidelity signal available to auditory cortex. These effects reveal a temporal processing deficit at multiple levels of the auditory pathway, thus reducing the fidelity of the auditory information available to higher-level regions necessary for speech perception and cognition. This impaired sensory encoding may contribute to known ELS-related problems with attention and learning.

### Neural loci of ELS effects on auditory gap detection

While there were deficits in temporal encoding at multiple levels of the auditory system, ELS-induced changes in ACx likely contributed to altered perception, because ACx combines inputs from the ascending auditory pathway and is necessary for gap detection. As we hypothesized, ELS during the critical period for ACx maturation has effects similar to those caused by auditory deprivation due to early hearing loss (Green et al., 2017). Conductive hearing loss in this period impairs temporal processing including perceptual and ACx gap detection, while also affecting ACx inhibitory kinetics, short-term plasticity, and synaptic strength (Sanes and Kotak, 2011; Caras and Sanes, 2015; Green et al., 2017). Because the perception of gaps relies specifically on activity of ACx inhibitory neurons (Weible et al., 2014), gap detection deficits in both hearing loss and ELS may arise from altered ACx inhibitory function. The ELS-induced reduction in non-monotonic ACx cells (Fig 6H) also points to inhibitory changes, as non-monotonicity in ACx is shaped by local inhibition (Wu et al., 2006). Similarly, altered inhibitory function in prefrontal cortex, hippocampus, and amygdala arising from ELS coincides with emotional and cognitive problems (Brenhouse and Andersen, 2011; Bath et al., 2016; Castillo-Gomez et al., 2017; Murthy et al., 2019; Page and Coutellier, 2019; Karst et al., 2020; Manzano Nieves et al., 2020). ELS causes early maturation of perineuronal nets, reduction in inhibitory neurons, and changes in BDNF levels in frontal cortex and amygdala (Cameron et al., 2017). These elements are involved in critical period regulation, including in ACx (Hensch, 2005; Do et al., 2015). Thus, the locus and details of ACx changes underlying ELS- and hearing loss-induced perceptual deficits may arise from common mechanisms involving critical period perturbation. An additional link is that both developmental stress and hearing loss reduce ACx excitability (Fig 6C,D) (Green et al., 2017). Since levels of cortical activity are tightly linked with mechanisms of plasticity, the overall reduction in neural activity by both manipulations indicates a common feature linking critical period dysregulation.

Although hearing loss and ELS cause similar changes to cortical responses, ELS caused peripheral changes not seen in animals with developmental hearing loss. Cochlear function is still immature during our early stress induction, suggesting that ELS could induce cochlear plasticity (Puel and Uziel, 1987; Arjmand et al., 1988; Mills and Rubel, 1996; Abdala and Keefe, 2012). Developmental conductive hearing loss does not affect the cochlea in terms of either frequency tuning (Caras and Sanes, 2015; Ye et al., 2021) or temporal processing (Yao and Sanes, 2018), measured as masked tuning curves and ABR thresholds to amplitude-modulated sounds, respectively. Yet, ELS impaired auditory nerve gap detection (Fig 4F,G, wave i). This could arise from direct effects of stress on elements of the cochlea. Stress is known to be otoprotective: acute stress induction protects against cochlear noise damage (Yoshida et al., 1999; Wang and Liberman, 2002). Studies have suggested that this protection arises from activation of glucocorticoid receptors within the cochlea (Jin et al., 2009), as well as from a local system that is molecularly equivalent to the HPA axis (Basappa et al., 2012). This raises the possibility that changes in glucocorticoid circulation could alter sound-evoked responses. For example, a mouse deficient in the corticotropin-releasing factor receptor displayed increased auditory sensitivity and greater susceptibility to noise damage (Basappa et al., 2012). This is a developmental effect which implicates systemic glucocorticoid effects on a cochlear stress axis as a signal to alter sound processing (Knipper et al., 2015). Yet here, wave i amplitude to only the 2^nd^ noise burst was reduced, indicating problems with recovery from adaptation rather than with auditory sensitivity, perhaps implicating involvement of vesicular release from inner hair cells.

Alternatively, stress may modulate efferent feedback from the olivocochlear system, which normally functions to refine inner hair cell sensitivity and frequency selectivity, and protect against hearing loss (Fuchs and Lauer, 2018). This efferent system is susceptible to top-down effects, as it can improve selective attention to sensory stimuli by modifying cochlear sensitivity (Delano et al., 2007; Gehmacher et al., 2022). Further, behaviorally-relevant plasticity has been demonstrated at wave i of the ABR (Rotondo and Bieszczad, 2020), and may indicate a role of olivocochlear feedback in perceptual learning (de Boer and Thornton, 2008; Irving et al., 2011; Fuchs and Lauer, 2018). Thus, ELS-induced changes in the brainstem, centrally, or directly on the cochlea could alter peripheral function.

Given that ELS resulted in both central and peripheral auditory effects, it is tempting to speculate about the relative contributions of peripheral and central gap detection deficits to perception. The impaired gap detection seen here in ACx neurons may be inherited from the poorer gap detection at the level of the auditory nerve. However, the ABR wave iv which reflects midbrain activity shows a recovery of gap detection encoding compared to earlier ABR waves from the auditory nerve and cochlear nucleus (Fig 4F,G). This may indicate some degree of central compensation, which is known to occur following hearing loss and aging (Caspary et al., 2008; Sanes and Kotak, 2011). The shapes of the response profiles for ELS and Controls across gap durations can be compared across behavioral, peripheral, and cortical responses, but yield an imperfect match. The response profiles for peripheral and cortical recordings were a general match for noise carriers (Fig 4F and 5K,L). Yet despite the behavior using a noise carrier, the final two behavioral sessions better matched the cortical tone carrier (Fig 3C and 5G,H). Further, within-animal measures of behavioral and ACx GDTs were not correlated for either tone or noise carriers. This is presumably due to the variability of the gap-PPI measure, the numbers of units recorded per animal, and the non-simultaneous measures, and can be addressed by future recordings conducted during gap-PPI sessions.

### Relations to known effects of ELS

While it is not surprising that altered HPA activity affects central auditory function, studies have consistently interpreted stress-induced auditory deficits in terms of altered sensory gating, a preattentional mechanism that filters irrelevant information by reducing the response to successive sounds (White and Yee, 1997; Rosburg et al., 2009). Experimentally, gating is a smaller P50 event-related potential response to the 2^nd^ (vs 1^st^) of two sounds separated by ∼500ms. Impaired gating is a biomarker for psychiatric disorders including PTSD and schizophrenia (Patterson et al., 2008; Javanbakht et al., 2011), and gating is altered by adult and early-life stress (White and Yee, 1997; Ellenbroek et al., 2004; Maxwell et al., 2006; Cromwell and Atchley, 2015; Ma et al., 2015; Yates et al., 2016). While sensory gating involves ACx (Josef-Golubic, 2020), gap detection operates on a much faster timescale – in fact, detection of gaps > 100ms does not require ACx, and thus sensory gating is likely probing a different phenomenon. Similarly, the behavioral phenomenon of sensorimotor gating (the reduction of a startle response by a preceding pulse) is also altered in stress and psychiatric disorders (Light and Braff, 1999; Ellenbroek et al., 2004; Huggenberger et al., 2013). However, sensorimotor gating is distinct from gap-PPI because it measures the response to a single sound, using pre-**pulse** inhibition (PPI) of acoustic startle. PPI does not require ACx, as it is intact with ACx inactivation (Threlkeld et al., 2008). Altered sensory gating is congruent with one of our results, because a reduced ratio in sensory gating can occur with a smaller P50 response to the 1^st^ of two sounds, which should predict smaller responses to single sounds. Indeed in rats, ELS-impaired gating is driven by a smaller P50 response to the first of two sounds (Ellenbroek et al., 2004), and this is consistent with our result of reduced firing of cortical neurons to simple tones (Fig 6C).

Consistent with our findings, there is some evidence that activation of the HPA axis has direct effects on auditory cortex. In human adults, corticosteroid treatment or stress exposure enhanced auditory evoked potentials and heightened auditory perceptual sensitivity (Ashton et al., 2000; Hasson et al., 2013). In adult rodents, application of a glucocorticoid agonist (dexamethasone) to ACx enhanced tone-evoked responses and broadened frequency tuning (Lei et al., 2014). Consistent with these results, adult-induced stress in rodents reduced dendritic length and complexity in the inferior colliculus and ACx, altered ACx inhibitory response properties, impaired auditory learning and attention, and enhanced ACx tone-evoked responses (Dagnino-Subiabre et al., 2005; Bose et al., 2010; Perez et al., 2013; Ma et al., 2015). While there is limited data on the direct effects of developmental stress on ACx, two studies in rats have shown that postnatal systemic dexamethasone administration or maternal separation increased synchrony to a 20 Hz stimulus in the left ACx, and reduced ACx evoked potential magnitudes (Ellenbroek et al., 2004; Yates et al., 2016).

One question is whether our effects are a result of top-down influences from regions known to be altered by ELS. For example, the amygdala has indirect effects on ACx via neuromodulatory inputs (Wenstrup et al., 2020), which may partially contribute to the reduced gap-PPI. Importantly, our neural recordings were conducted under anesthesia, minimizing the impact of top-down influences from higher-level regions (Mashour, 2014; Cai et al., 2016). Further, we used a short (50ms) interval between stimulus and startle in our gap-PPI perceptual measure to ensure that it was pre-attentive (Li et al., 2009), reducing the impact of higher-level regions.

In conclusion, the behavioral deficits seen here may arise from plasticity at both the level of ACx and earlier in the auditory pathway. Critical period mechanisms can now be examined to identify elements that are altered by early-life stress, and whether those differ from elements altered by disruptions in early auditory experience, such as developmental hearing loss.

## Author Contributions

M.J.R designed the research concepts and the experiments; Y.Y. and M.M. collected the data; M.J.S. and J.D.G processed the data; Y.Y. and M.J.R. analyzed the data; M.J.R. wrote the manuscript.

## Acknowledgements

This work was supported by NIDCD R01 DC013314 to M.J.R. The content is solely the responsibility of the authors and does not necessarily represent the official views of the National Institutes of Health. The authors would like to thank Dr. Julia Huyck for comments on an earlier version of the manuscript.

## Data availability

The data are available from the corresponding author on reasonable request.

